# Development of a Temperature-Based Method for Repeated Markerless Gene Integration in cyanobacterium *Synechococcus elongatus* PCC 7942

**DOI:** 10.64898/2026.11.15.766822

**Authors:** Kazuma Ohdate, Taro Tsuboi, Hazuki Hasegawa, Kan Tanaka, Satoru Watanabe

## Abstract

Genetic engineering of cyanobacteria can require multiple rounds of genome modification, highlighting the need for flexible markerless engineering tools. Here, we established a temperature-based markerless gene integration method using the temperature-sensitive H1 strain of *Synechococcus elongatus* PCC 7942, which, unlike the wild-type strain, cannot grow at 40 °C. Disruption of *nynA*, encoding a putative endoribonuclease, allowed H1 to grow at 40 °C, enabling transformants with genes integrated at the *nynA* locus to be selected without an antibiotic-resistance marker. Complementation of *nynA* at a separate genomic locus restored temperature sensitivity, allowing the same selection principle to be reused. Using this strategy, *gfp*, *mScarlet*, and *luciferase* were sequentially integrated, with all three genes retained and expressed. To demonstrate its applicability to metabolic engineering, markerless integration of *cnsA*, encoding cineole synthase, enabled 1,8-cineole production at 81.4 ± 9.52 µg g⁻¹ wet cell weight. This strategy enables repeated markerless genome modification in the H1 strain using temperature-based selection.

## INTRODUCTION

Cyanobacteria have attracted attention as promising platforms for sustainable bioproduction because they can directly convert CO₂ into valuable compounds using light energy through photosynthesis ^1^. Recent advances in cyanobacterial genetic engineering, including inducible expression systems ^2, 3^ and autonomously replicating vectors ^4, 5^, have facilitated the production of various useful compounds, including biofuel precursors and high-value compounds ^6^. Combining these approaches with strategies such as gene knockout and knockdown has further improved production yields ^7, 8^. As these engineering strategies become increasingly complex, stepwise modification of multiple genes is often required. Therefore, tools that enable flexible and repeated genome modification are increasingly important for cyanobacterial engineering ^9^.

Positive selection using antibiotic-resistance genes is widely used to isolate bacterial transformants. However, repeated genetic modification can lead to the accumulation of antibiotic-resistance markers in the genome, limiting the availability of selection markers for subsequent modifications. Markerless genetic engineering methods have therefore been developed to remove or avoid retaining selection markers after genome modification ^10^. By allowing selection markers to be reused, these methods facilitate iterative genome engineering without the accumulation of antibiotic-resistance markers ^11^.

In cyanobacteria, counterselection markers such as the endoribonuclease MazF and the levansucrase SacB have been used for markerless genetic engineering, with such systems demonstrated in species including *Synechocystis* sp. PCC 6803, *Synechococcus elongatus* PCC 7942 (*Synechococcus* 7942), and *Anabaena* sp. PCC 7120 ^12, 13, 14^. These approaches demonstrate the utility of counterselection for markerless genome modification. However, expanding the range of available selection principles would be useful for cyanobacterial genetic engineering, particularly for applications requiring multiple rounds of genetic modification.

We focused on the temperature-sensitive H1 strain of *Synechococcus* 7942, which exhibits severely impaired growth at high temperature. The H1 strain carries a G60T mutation in the gene encoding tRNA-Leu (CAA), and genetic analysis has shown that disruption of *nynA*, which encodes a putative endoribonuclease, enables growth at 40 °C^15^. Although the mechanism by which *nynA* disruption abolishes high-temperature sensitivity remains unclear, we hypothesized that the relationship between *nynA* disruption and the temperature-dependent growth phenotype could be exploited as a selectable phenotype. Specifically, because disruption of *nynA* enables the H1 strain to grow at 40 °C, homologous recombinants targeting *nynA* could potentially be selected by incubation at 40 °C without using an antibiotic-resistance marker.

In this study, we established a markerless gene integration method in the H1 strain by exploiting its temperature sensitivity and *nynA*, using incubation at 40 °C as the selection pressure. Furthermore, complementation of *nynA* at a separate genomic locus restored the temperature-sensitive phenotype, enabling multiple rounds of gene integration using the same selection principle (Fig. 1). We demonstrated the utility of this method for both reporter gene introduction and metabolic engineering by applying it to 1,8-cineole production ^16, 17^. These results establish a temperature-based strategy for repeated markerless genome modification in the H1 strain.

**Figure 1.**
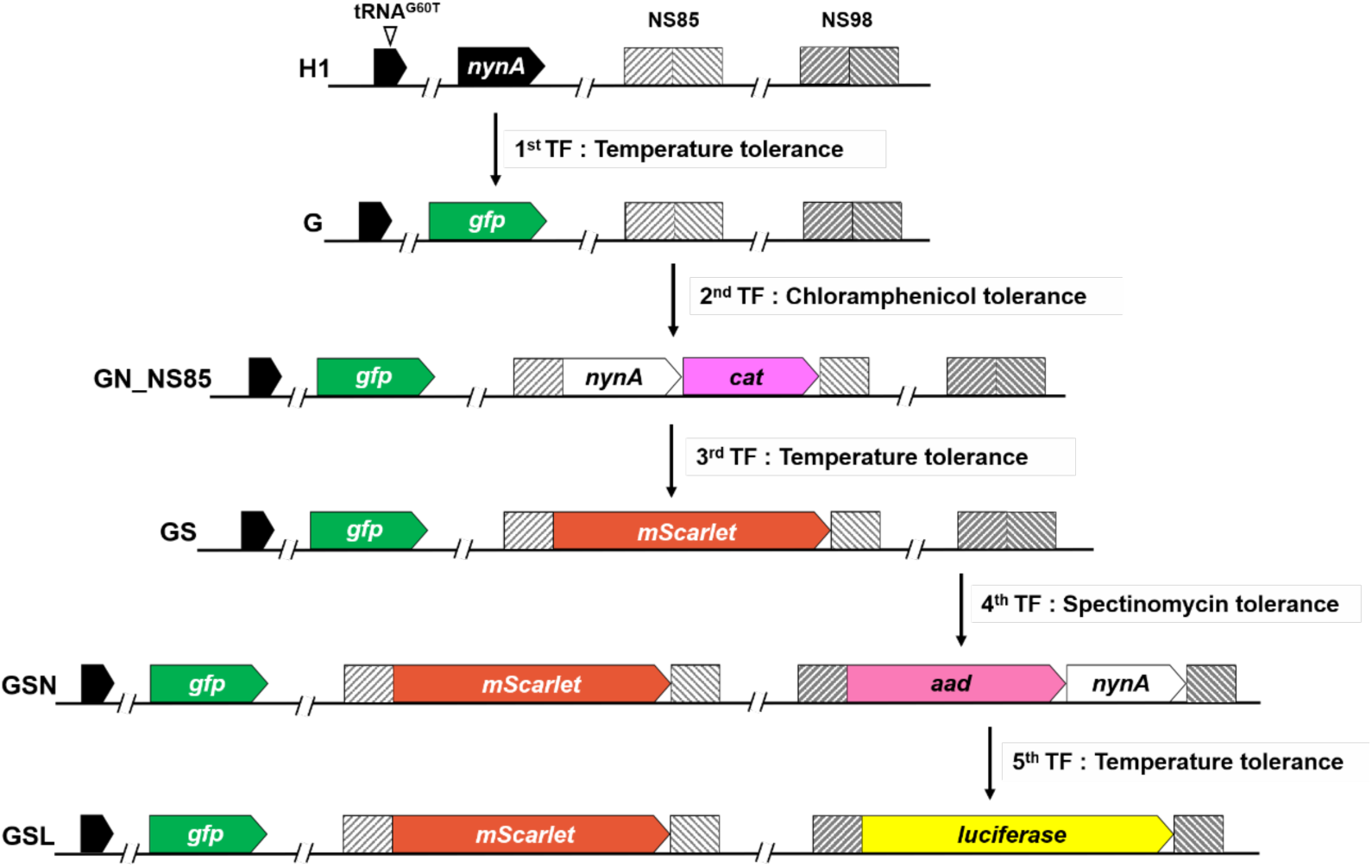
Schematic overview of the markerless transformation strategy using temperature-based selection. Sequential transformation steps and the resulting strains are shown.

## RESULTS

### High-temperature selection enables isolation of transformants in the H1 strain

We first tested whether the temperature-sensitive phenotype of the H1 strain could be exploited for high-temperature selection of transformants. Because disruption of *nynA* restores growth of the H1 strain at 40 °C, we generated the integration construct p1640_aad to introduce the spectinomycin-resistance gene *aad* into the *nynA* locus (Fig. 2A). Transformants were selected using spectinomycin or by incubation at 40 °C (Fig. 2B). Selection by incubation at 40 °C yielded significantly more colonies than selection with spectinomycin (Fig. S1A). We then randomly picked 100 colonies obtained under 40 °C selection and examined their ability to grow at 40 °C and on spectinomycin-containing medium (Figs. 2C and S1B). All 100 colonies exhibited both high-temperature tolerance and spectinomycin resistance, suggesting a low frequency of false-positive colonies under high-temperature selection. Furthermore, PCR analysis confirmed integration of *aad* at the targeted *nynA* locus in transformants obtained under both selection conditions. Complete segregation of the *aad*-containing genome was confirmed in transformants obtained by incubation at 40 °C (Fig. 2D). These results demonstrate that transformants carrying genes integrated at the *nynA* locus can be selected by incubation at 40 °C, providing a basis for markerless gene integration in the H1 strain.

**Figure 2.**
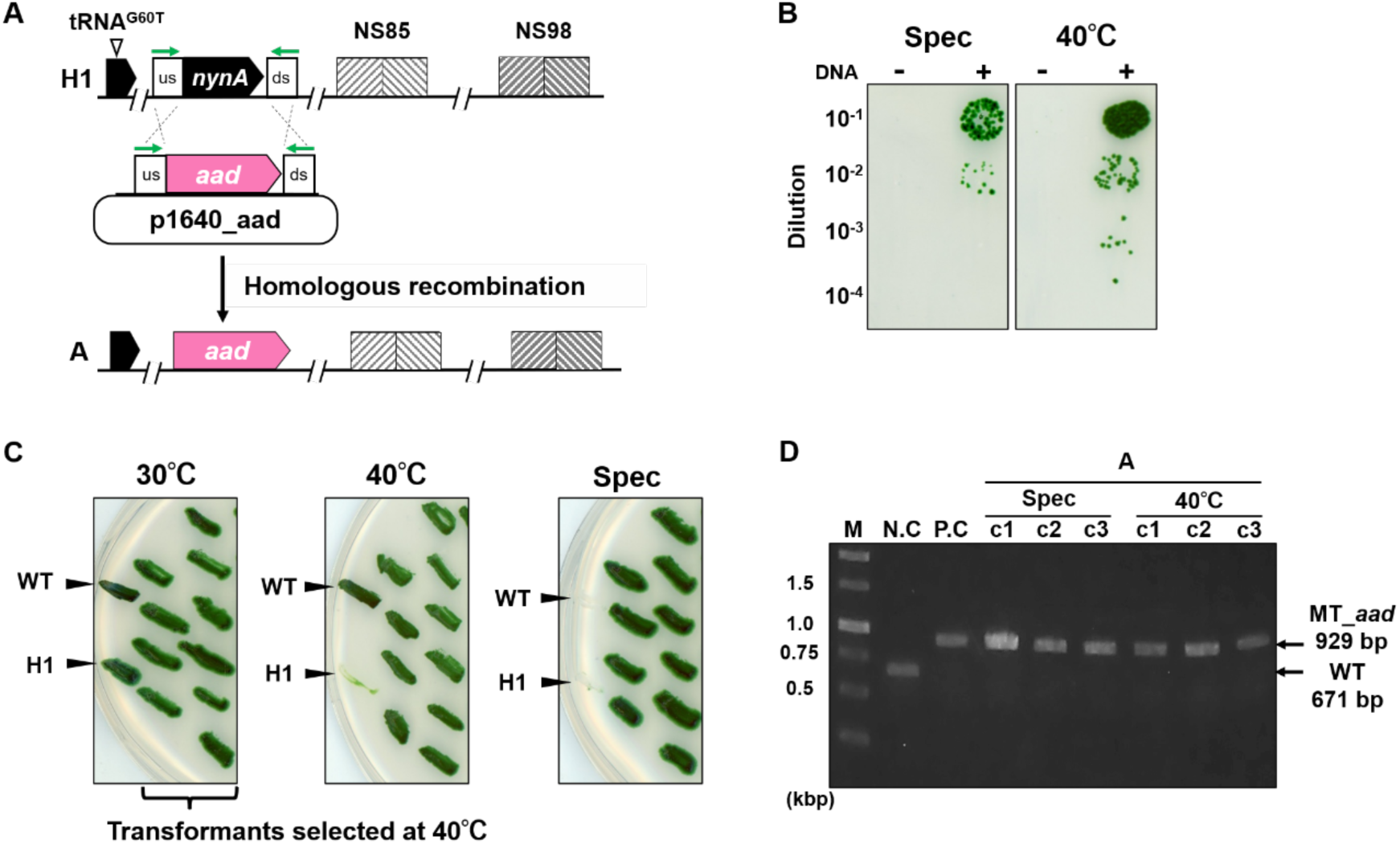
Validation of temperature-based selection using an *aad* marker. (A) Schematic representation of the integration construct used to introduce *aad* into the H1 strain. Green arrows indicate the positions of primers used for PCR analysis. (B) Selection of transformants with spectinomycin or incubation at 40 °C. Transformants were diluted and spotted onto spectinomycin-containing BG-11 agar plates and incubated at 30 °C onto antibiotic-free BG-11 agar plates and incubated at 40 °C for 7 days. (C) Phenotypic characterization of transformants obtained by incubation at 40 °C. One hundred transformants were tested for temperature and spectinomycin resistance under the indicated conditions. A representative portion of the plates showing 10 of the 100 tested transformants is presented; the complete results for all 100 transformants are shown in Fig. S1B. WT and H1 strains were included as controls. (D) PCR analysis of H1 and independent transformants obtained by selection with spectinomycin or incubation at 40 °C. Primers shown as green arrows in Fig. 2A were used. M, DNA marker; NC, H1 strain; PC, p1640_aad plasmid (Table S1).

### Markerless gene integration using high-temperature selection

To determine whether the temperature-based selection could be used for markerless gene integration, we attempted to introduce a *gfp* expression construct without an antibiotic-resistance marker into the H1 strain. We generated the integration construct p1640_gfp to introduce *gfp* into the *nynA* locus (Fig. 3A). Following transformation, transformants were selected by incubation at 40 °C (Fig. S2A). PCR analysis confirmed integration of the *gfp* expression construct at the targeted locus (Fig. 3B). The resulting *gfp*-integrated strain grew at 40 °C, like the wild-type strain, whereas the parental H1 strain did not (Fig. 3C). GFP expression was subsequently evaluated by flow cytometry (Fig. 3D) and fluorescence microscopy (Fig. S2B). GFP fluorescence was detected by both methods, confirming functional expression of the integrated *gfp* gene. These results demonstrate that markerless gene integration can be achieved in the H1 strain by targeting *nynA* and selecting at 40 °C. However, disruption of *nynA* renders the resulting strain tolerant to 40 °C, preventing the same temperature-based selection from being directly reused for a subsequent round of markerless gene integration.

**Figure 3.**
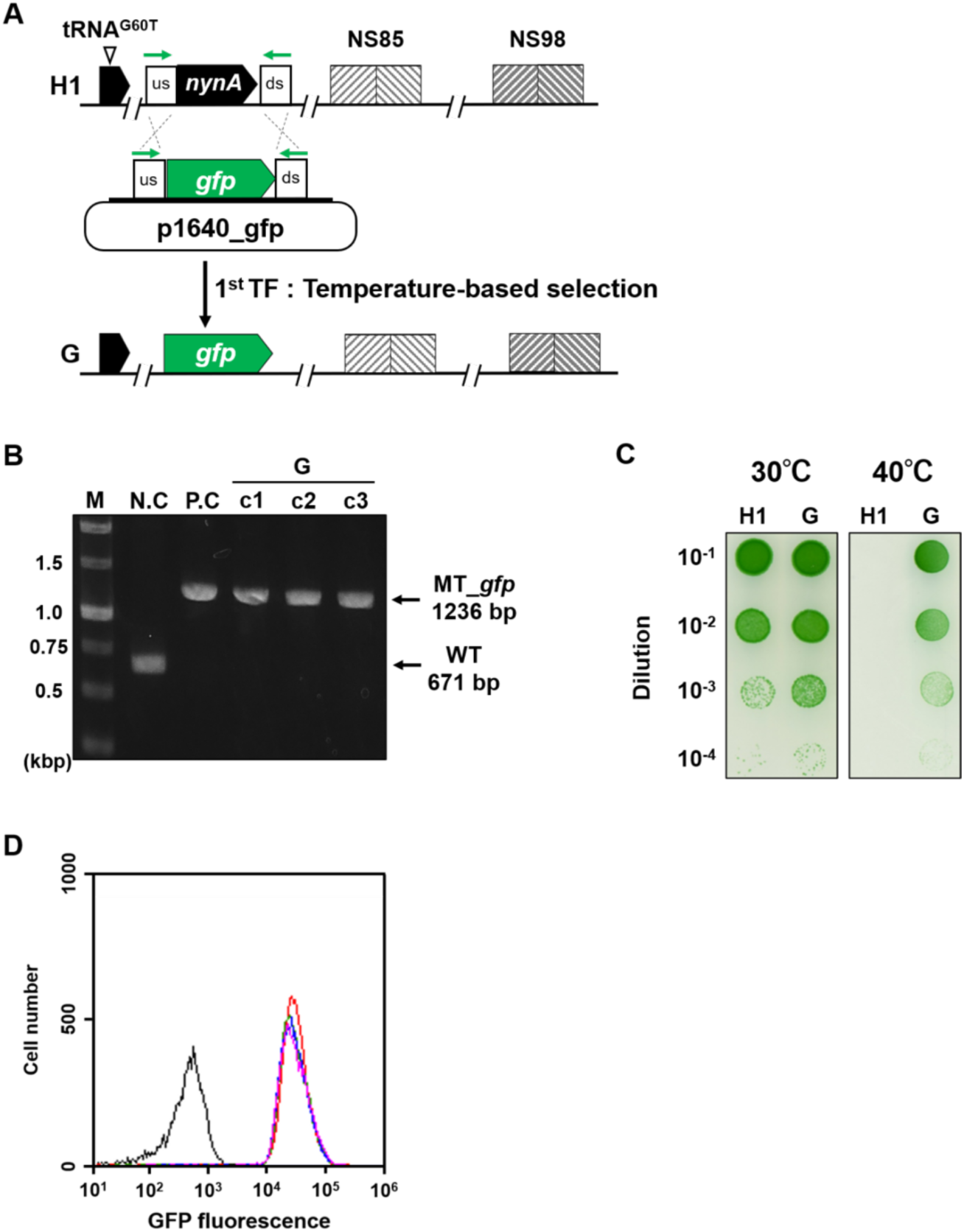
Construction and characterization of the *gfp*-expressing strain using temperature-based selection. (A) Schematic representation of the integration construct used to introduce the *gfp* expression construct into the *nynA* locus of H1 strain. Green arrows indicate the positions of primers used for PCR analysis. (B) PCR analysis of H1 and three independent transformants (c1–c3). Primers shown as green arrows in Fig. 3A were used to confirm integration of the *gfp* gene. M, DNA marker; NC, H1 strain; PC, p1640_gfp plasmid (Table S1). (C) Growth of H1 and G strains at 30 and 40 °C. Strains were diluted and spotted onto BG-11 agar plates and incubated at 30 or 40 °C for 5 days. (D) Flow cytometric analysis of GFP fluorescence. Histograms of GFP fluorescence are shown for the H1 strain (black), NS-GFP (red) as a GFP-positive control, and G strains c1 (green), c2 (blue), and c3 (pink).

### Restoration of temperature sensitivity by *nynA* complementation

We next investigated whether complementation of *nynA* at a separate genomic locus in the *gfp*-integrated strain could restore the temperature-sensitive phenotype. We generated the integration construct pNS85_nynA to introduce *nynA* together with the chloramphenicol-resistance gene *cat* into the NS85 locus (Fig. 4A). Transformants were selected on BG-11 plates containing chloramphenicol, and PCR analysis confirmed integration of *nynA* and *cat* at the targeted NS85 locus (Fig. S3A). The resulting GN strain was unable to grow at 40 °C, like the parental H1 strain, indicating restoration of the temperature-sensitive phenotype (Fig. 4B). Thus, *nynA* complementation restored the temperature-sensitive phenotype required for another round of high-temperature selection.

**Figure 4.**
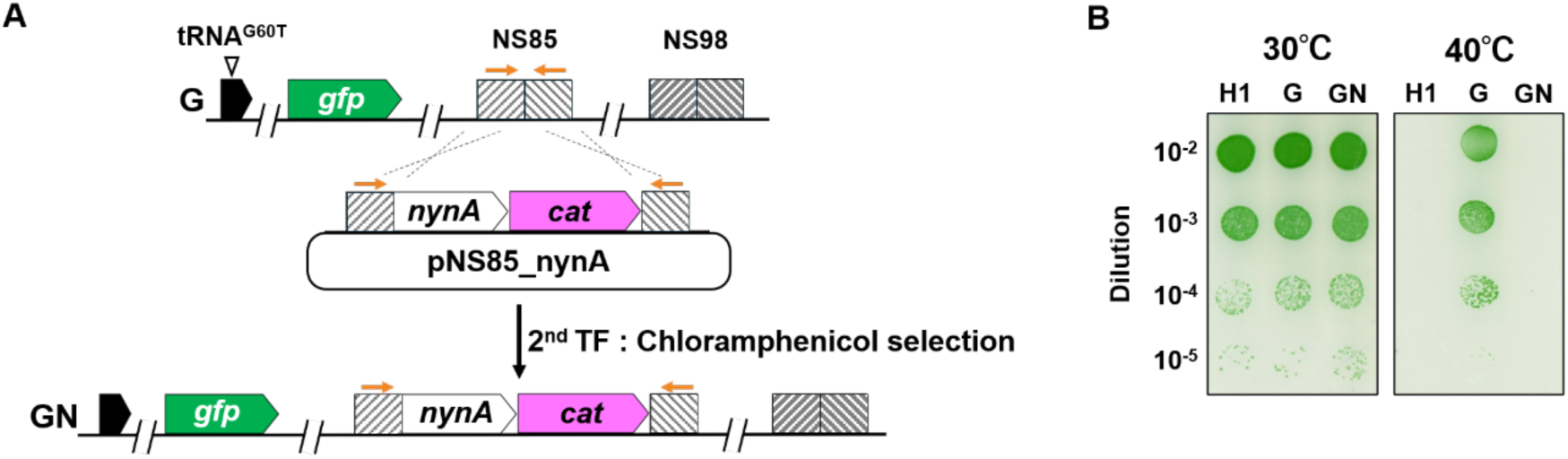
Construction and characterization of the *nynA*-complemented strain. (A) Schematic representation of the integration construct used to introduce *nynA* into the NS85 locus of the G strain. Orange arrows indicate the positions of primers used for PCR analysis. (B) Temperature-sensitive phenotype of the GN strain. H1, G, and GN strains were diluted and spotted onto BG-11 agar plates and incubated at 30 °C or 40 °C for 5 days.

### Second round of markerless gene integration

We next tested whether this restored temperature sensitivity could be used for a second round of markerless gene integration. Using the temperature-sensitive GN strain, we introduced pNS85_mScarlet, an integration construct carrying *mScarlet* without an antibiotic-resistance marker, into the locus containing the complemented *nynA* and *cat* genes (Fig. 5A). Transformants were selected by incubation at 40 °C (Fig. S4A), and PCR analysis confirmed integration of the *mScarlet* expression construct at the targeted locus (Fig. 5B). The resulting GS strain grew at 40 °C and was sensitive to chloramphenicol, consistent with replacement of the complemented *nynA* and *cat* genes by the *mScarlet* expression construct (Fig. 5C). mScarlet fluorescence was observed by both flow cytometry and fluorescence microscopy (Figs. 5D and S4B). Importantly, GFP fluorescence was also detected in the GS strain (Figs. 5D and S4B). PCR analysis confirmed that the previously integrated *gfp* gene remained intact in the genome (Fig. S4C). These results demonstrate that markerless gene integration can be repeated by restoring the temperature-sensitive phenotype through *nynA* complementation at a new genomic locus and subsequently targeting the complemented *nynA* locus for markerless integration.

**Figure 5.**
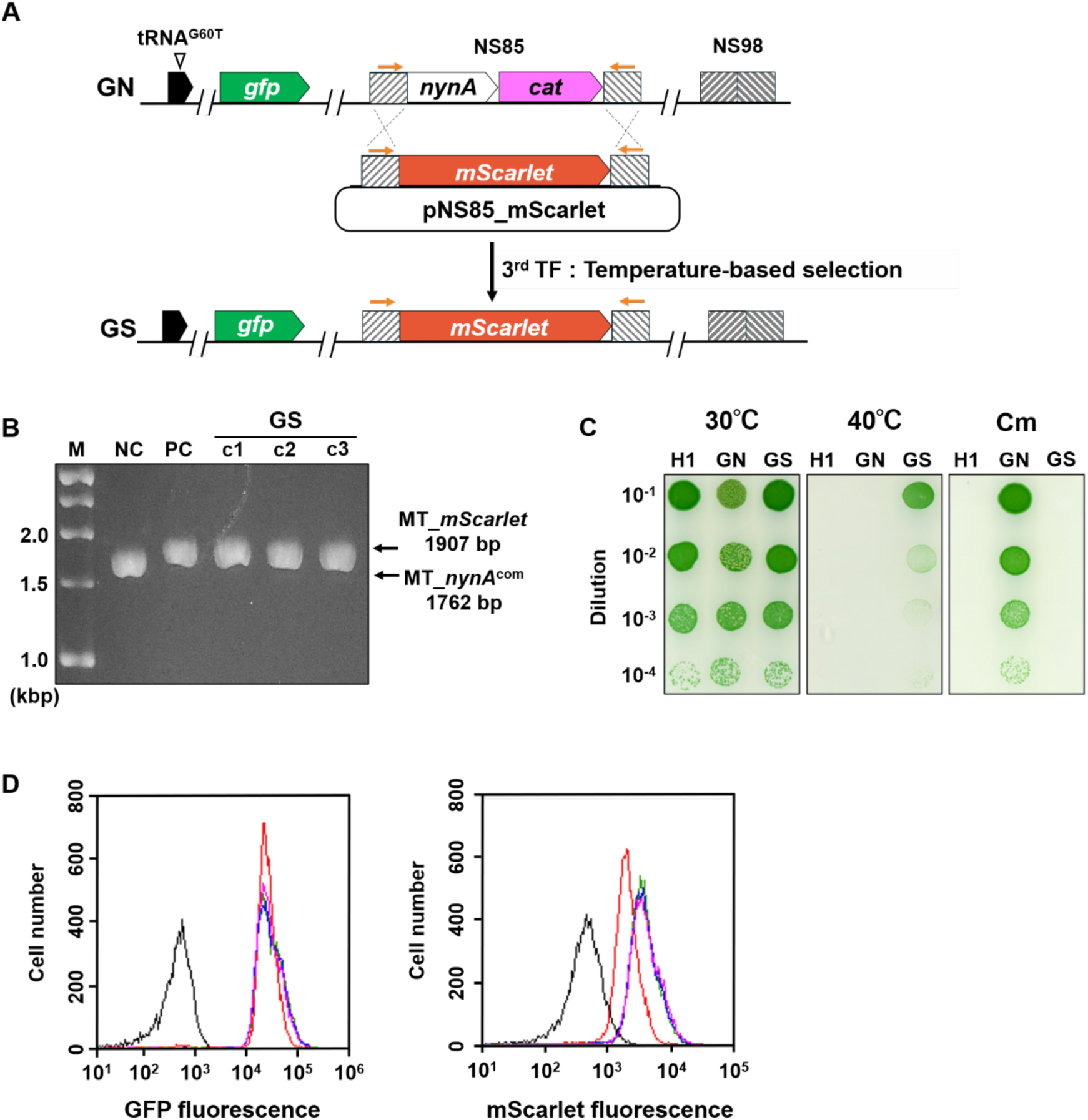
Construction and characterization of the *mScarlet*-expressing strain using temperature-based selection. (A) Schematic representation of the integration construct used to introduce the *mScarlet* expression construct into the complemented *nynA* locus of the GN strain. Orange arrows indicate the positions of primers used for PCR analysis. (B) PCR analysis of the GN strain and three independent transformants (c1–c3). Primers shown as orange arrows in Fig. 5A were used to confirm integration of the *mScarlet* gene. M, DNA marker; NC, GN strain; PC, pNS85_mScarlet plasmid. PCR products were separated on a 1.8% agarose gel in TBE buffer at 50 V. (C) Growth phenotype of the GS strain under selection and nonselection conditions. H1, GN, and GS strains were diluted and spotted onto BG-11 agar plates with or without chloramphenicol and incubated at 30 or 40 °C for 5 days. (D) Flow cytometric analysis of GFP and mScarlet fluorescence. Histograms of GFP and mScarlet fluorescence are shown for the H1 strain (black), GN strain (red), and GS strains c1 (green), c2 (blue), and c3 (pink).

### Third round of markerless gene integration by repeated *nynA* complementation

To test whether markerless gene integration could be extended to a third round, *nynA* was complemented at the NS98 locus together with the spectinomycin-resistance gene *aad*. The resulting GSN strain exhibited the temperature-sensitive phenotype required for the next round of selection (Figs. 6A, B, S3B). An integration construct carrying *luciferase* gene without an antibiotic-resistance marker was then introduced into the locus containing the complemented *nynA* and *aad* genes. Transformants were selected by incubation at 40 °C (Figs. 6A, C, and S5A). The resulting GSL strain grew at 40 °C and was sensitive to spectinomycin, consistent with replacement of the complemented *nynA* and *aad* genes by the *luciferase* expression construct (Fig. 6D). Luciferase expression was confirmed by a luciferase assay (Fig. 6E). GFP and mScarlet fluorescence were retained in the resulting strain, and PCR analysis confirmed that the corresponding genes remained intact in the genome (Figs. S5B-D). Thus, three heterologous genes, *gfp*, *mScarlet*, and *luciferase*, were sequentially integrated into the genome without an antibiotic-resistance marker. These results demonstrate that repeated *nynA* complementation enables successive rounds of markerless gene integration while preserving previously introduced genes.

**Figure 6.**
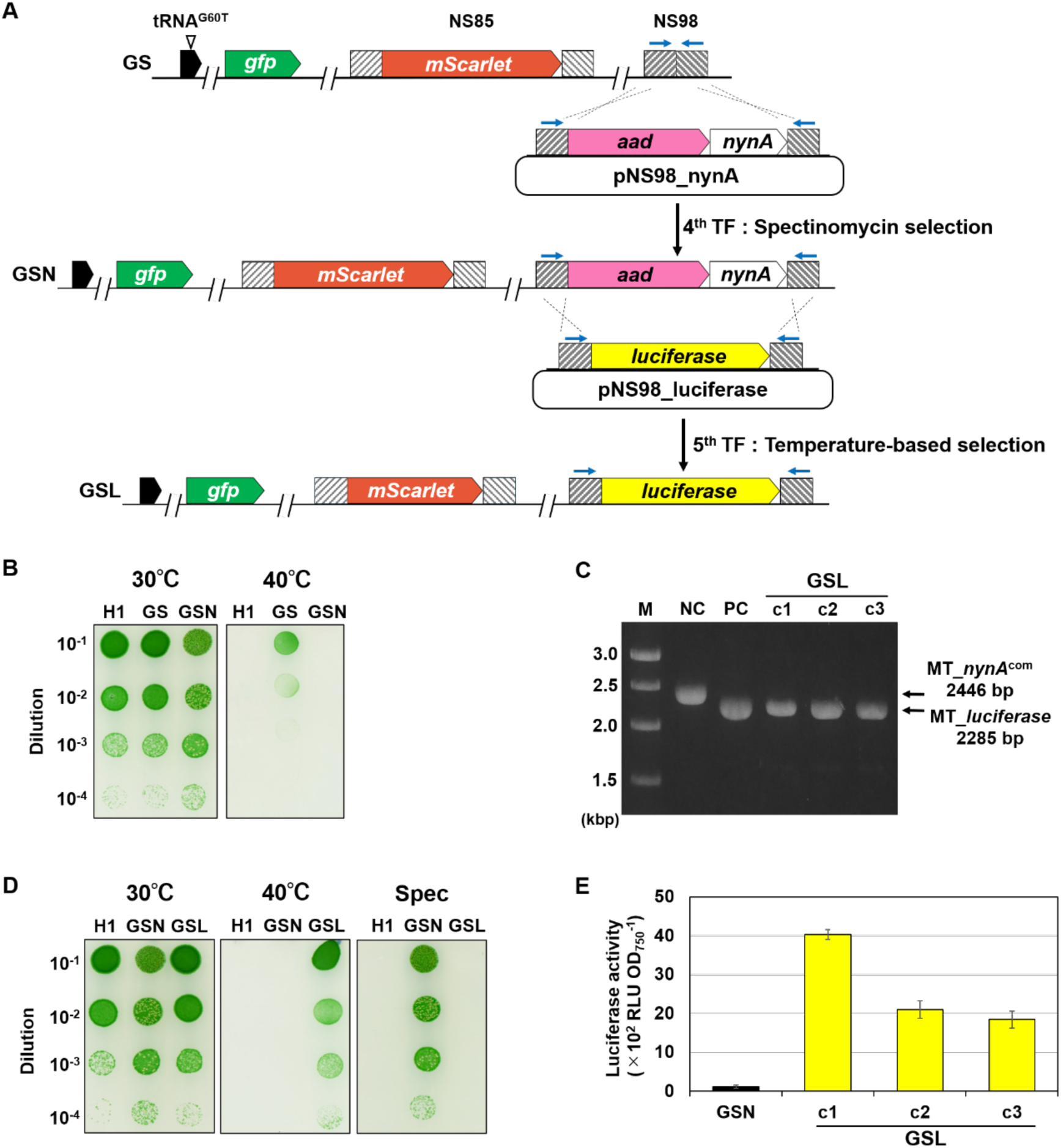
Construction and characterization of the *luciferase*-expressing strain by repeated *nynA* complementation and temperature-based selection. (A) Schematic representations of the integration constructs used to introduce *nynA* into the NS98 locus of the GS strain and the *luciferase* expression construct into the complemented *nynA* locus of the GSN strain. Blue arrows indicate the positions of primers used for PCR analysis. (B) Growth of H1, GS, and GSN strains at 30 and 40 °C. Strains were diluted and spotted onto BG-11 agar plates and incubated at 30 or 40 °C for 5 days. (C) PCR analysis of the GSN strain and three independent transformants (c1–c3). Primers shown as blue arrows in Fig. 6A were used to confirm integration of the *luciferase* expression construct. M, DNA marker; NC, GSN strain; PC, pNS98_luciferase plasmid. PCR products were separated on a 1.5% agarose gel in TBE buffer at 50 V. (D) Growth of H1, GSN, and GSL strains at 30 and 40 °C with or without spectinomycin. Strains were diluted and spotted onto BG-11 agar plates and incubated for 5 days. (E) Luciferase activity of the GSN and GSL strains. Relative Luciferase activity was calculated as RLU OD_750_^-1^. Data is presented as the mean ± SD of three biological replicates. Experimental details are described in the Materials and Methods.

### Application of markerless gene integration to 1,8-cineole production

To test the utility of this strategy in metabolic engineering, we utilized the *cnsA* gene, which encodes cineole synthase (CnsA). CnsA catalyzes the conversion of geranyl pyrophosphate (GPP) to 1,8-cineole through a series of reactions ^16^. The integration construct p1640_cnsA, carrying *cnsA* without an antibiotic-resistance marker, was introduced into the H1 strain, and transformants were selected by incubation at 40 °C (Fig. 7A). The *cnsA* gene was placed under the control of a LacI-regulated promoter. PCR analysis confirmed integration of the *cnsA* expression construct at the *nynA* locus, and the resulting strain exhibited the expected high-temperature-tolerant phenotype (Figs. 7B and 7C). We next evaluated whether the integrated *cnsA* gene was functional by inducing its expression with isopropyl ß-D-1-thiogalactopyranoside (IPTG) and analyzing 1,8-cineole production by gas chromatography (Fig. 7D). The *cnsA*-integrated strain produced 1,8-cineole at 81.4 ± 9.52 µg g⁻¹ wet cell weight (WCW), whereas no corresponding signal was detected in the H1 strain. This production level was comparable to that previously reported for a CnsA-expressing strain ^17^. Thus, *cnsA* was introduced and functionally expressed without an antibiotic-resistance marker, resulting in production of the target compound. Together with the sequential integration experiments described above, these results demonstrate the applicability of this temperature-based strategy to markerless gene integration for metabolic engineering.

**Figure 7.**
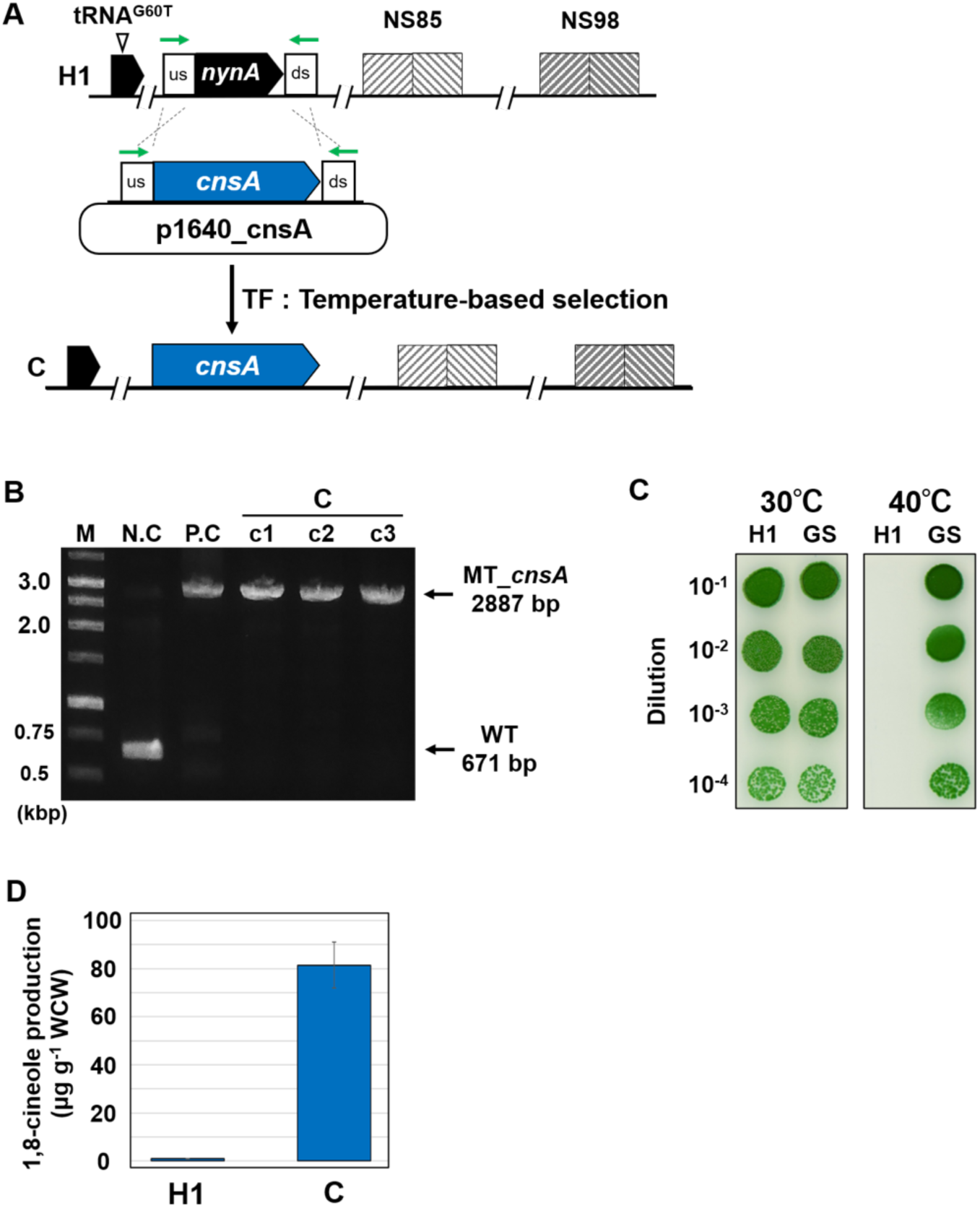
Construction and characterization of the *cnsA*-expressing strain using temperature-based selection. (A) Schematic representation of the integration construct used to introduce the *cnsA* expression construct into the *nynA* locus of H1 strain. Green arrows indicate the positions of primers used for PCR analysis. (B) PCR analysis of H1 and three independent transformants (c1–c3). Primers shown as green arrows in Fig. 7A were used to confirm integration of the *cnsA* expression construct. M, DNA marker; NC, H1 strain; PC, p1640_cnsA plasmid (Table S1). (C) Growth of H1 and C strains at 30 and 40 °C. Strains were diluted and spotted onto BG-11 agar plates and incubated at 30 or 40 °C for 5 days. (D) 1,8-cineole production by H1 and C strains. Production levels were normalized to wet cell weight (WCW). Data is presented as the mean ± SD of three biological replicates.

## DISCUSSION

In this study, we established a markerless gene integration method in the temperature-sensitive H1 strain of *Synechococcus* 7942 by combining gene integration at the *nynA* locus with selection by incubation at 40 °C. Furthermore, complementation of *nynA* at a separate genomic locus restored the temperature-sensitive phenotype, enabling repeated rounds of markerless gene integration using the same selection principle. Indeed, three heterologous genes, *gfp*, *mScarlet*, and *luciferase*, were sequentially integrated into the genome without retaining an antibiotic-resistance marker, and their retention and expression were confirmed. The utility of this approach was further demonstrated by the introduction of *cnsA*, which the production of enabled production of 1,8-cineole, a monoterpenoid with reported anti-inflammatory properties ^18^, extending its application beyond reporter gene introduction to metabolic engineering.

In cyanobacteria, counterselection methods based on SacB or MazF have previously been developed for markerless genome modification ^12, 13, 14^. Unlike these approaches, our method uses the temperature-dependent growth phenotype of the H1 strain as the basis for selection. Importantly, *nynA* complementation restores temperature sensitivity, allowing the same selection principle to be reused for successive rounds of gene integration. This strategy enables repeated genome modification without the accumulation of antibiotic-resistance markers.

Although *nynA* is predicted to function as an RNase ^15^, its enzymatic activity has not been experimentally confirmed, and the mechanism by which *nynA* disruption abolishes high-temperature sensitivity remains unclear. Changes in RNase function have been reported to affect temperature-dependent growth and stress responses in other bacteria. For example, in *E. coli*, inactivation of RNase III has been reported to affect the expression of various genes and reduce thermotolerance ^19^. These findings raise the possibility that NynA may influence high-temperature sensitivity indirectly through changes in cellular processes associated with temperature-dependent growth or stress responses.

Elucidating how disruption of *nynA* abolishes high-temperature sensitivity may provide insights into the development of similar temperature-based selection systems in other microorganisms. At present, the applicability of this method has been demonstrated only in the H1 strain of *Synechococcus* 7942. Therefore, evaluating its applicability to other cyanobacteria will be an important direction for future studies. Further optimization of *nynA* complementation, including the choice of genomic integration site and promoter strength, may improve the efficiency and stringency of high-temperature selection.

In conclusion, we established a markerless gene integration method that enables repeated gene integration by combining the high-temperature sensitivity of the H1 strain with *nynA* complementation. This method provides an additional strategy for multistep markerless genome modification in the H1 strain, with potential applications in cyanobacterial metabolic engineering.

## MATERIALS AND METHODS

### Cyanobacteria growth condition

The freshwater cyanobacterium *Synechococcus* 7942 H1 strain, a temperature-sensitive mutant carrying a G60T substitution in tRNA-Leu (CAA) ^15^, was used as the host strain in this study. Cells were cultured in modified BG-11 medium supplemented with 35.3 mM NaNO₃ (twice the standard BG-11 concentration) and 20 mM HEPES-KOH (pH 8.0). Cultures were grown photoautotrophically at 30 °C under continuous illumination (50 μmol photons m⁻² s⁻¹). Unless otherwise stated, cultures (20 mL) were grown in 100 mL Erlenmeyer flasks with shaking at 140 rpm. When required, spectinomycin (40 μg mL⁻¹) or chloramphenicol (7.5 μg mL⁻¹) was added to the medium.

### Plasmid and strain construction

DNA fragments were amplified using KOD One DNA polymerase (TOYOBO, Osaka, Japan) and subcloned into vector plasmids using the In-Fusion HD Cloning Kit (TaKaRa Bio, Shiga, Japan). The *nynA* and *aad* genes were amplified from the genomic DNA of the HDN-1 strain ^15^ using a specific primer set (F1/R2). The amplified fragments were cloned into the pUC19 vector, which had been linearized by polymerase chain reaction (PCR) using the primer set (R3/F4), generating the plasmid p1640_aad. To generate the plasmids p1640_gfp and p1640_cnsA, the respective insertion fragments were amplified from the pYS-1C-GFP and pYS-cnsA plasmids ^4, 17^ using specific primer sets (F5/R6 for p1640_gfp, F7/R8 for p1640_cnsA). The *cnsA* sequence used in p1640_cnsA was codon-optimized for expression in *Synechococcus* 7942. The amplified fragments were then cloned into the p1640 vector, which had been linearized using the primer set (F9/R10). These plasmids were introduced into the H1 strain. Following incubation at 40°C, transformants exhibiting a suppressed temperature-sensitive phenotype were obtained. Successful transformation of the strains was confirmed by PCR amplification using a specific primer set (F26/R27). To generate the plasmids pNS85_nynA and pNS85_mScarlet, the respective insertion fragments were amplified from *Synechococcus* 7942 genomic DNA and the pEX2-mScarlet plasmid ^20^ using specific primer sets (F11/R12 for pNS85_nynA, F13/R14 for pNS85_mScarlet). The amplified fragments were cloned into the pNS85 vector, which had been linearized using the primer sets (F15/R16 for pNS85_nynA, F17/R16 for pNS85_mScarlet). These plasmids were introduced into the G strain. Successful transformation of the strains was confirmed by PCR amplification using specific primer sets (F28/R29). To generate the plasmids pNS98_nynA and pNS98_luciferase, the respective insertion fragments were amplified from *Synechococcus* 7942 genomic DNA and the pEYN12 plasmid ^3^ using specific primer sets (F18/R19 for pNS98_nynA, F20/R21 for pNS98_luciferase). The amplified fragments were cloned into the pNS98 vector, which had been linearized using the primer sets (F22/R23 for pNS98_nynA, F24/R25 for pNS98_luciferase). These plasmids were introduced into the GS strain. Successful transformation of the strains was confirmed by PCR amplification using specific primer sets (F30/R31).

### Microscopy

Fluorescent images were obtained using BX53 microscope (OLYMPUS, Tokyo, Japan) at ×40 or 100 magnification with a DP71 digital camera (OLYMPUS) and DP Controller software ver. 3. 3. 1. 292 (OLYMPUS).

### Flow cytometry

Cells were analyzed by fluorescence-activated cell sorting (FACS) based on GFP and mScarlet fluorescence using a BD Accuri™ C6 Flow Cytometer (BD Biosciences, San Jose, CA, USA) and BD CFlow software (BD Biosciences), as previously described ^21^. Cyanobacterial cells were identified based on chlorophyll autofluorescence detected on the FL3 channel. GFP and mScarlet fluorescence were measured using the FL1 and FL2 channels, respectively. The *Synechococcus* 7942 NS-GFP strain was included as a GFP-positive control.

### Induction of Luciferase and Luciferase Assay

Luciferase activity was measured essentially as described previously ^3^, using the Luciferase Assay System (Promega, Madison, WI, USA), except that luminescence was recorded with a Nivo multimode plate reader (PerkinElmer, Shelton, CT, USA). Briefly, luciferase expression was induced with theophylline (2 mM final concentration). Cells were then harvested by centrifugation and lysed by freeze–thaw treatment followed by incubation in lysis buffer. Cell lysates were mixed with Luciferase Assay Reagent, and luminescence was measured for 5 s. Relative luciferase activity was normalized to OD_750_ and expressed as RLU OD ^-1^.

### Induction of CnsA and recovery of 1,8-cineole as a volatile component

Production of 1,8-cineole was analyzed as previously described ^17^. Briefly, *cnsA* expression was induced with IPTG (1 mM final concentration), and volatile compounds in the headspace were collected using a SPME fiber (50/30 μm PDMS/DVB/CAR; Supelco, Bellefonte, PA, USA). The collected volatiles were analyzed by gas chromatography as described previously.

## ASSOCIATED CONTENT

The lists of stain, plasmid, and primers (Table S1-S3) and Supplementary figures (Fig. S1-S5)

## AUTHOR INFORMATION

### Author contributions

H.H. K.T. and S.W. designed the concept and the experiments of this study; K.O., T.T. and S.W. performed the experiments; K.O., and S.W. analyzed the data; K.O., T.K., and S.W. wrote the manuscript.

## ACKNOWLEDGMENTS

We are grateful to Prof. Taku Chibazakura and Prof. Kei Asai for their valuable comments on the concept of this study. This work was supported by the Ministry of Education, Culture, Sports, Science and Technology of Japan to SW (23H02130, 24H00869, and 24H00871) and New Energy and Industrial Technology Development Organization (NEDO, JPNP17005) (to S.W.).

## ABBREVIATIONS

*Synechococcus* 7942: *Synechococcus elongatus* PCC 7942
WT: Wild-type
WCW: wet cell weight
*E*. *coli*: *Escherichia coli*
IPTG: isopropyl ß-D-1-thiogalactopyranoside

## REFERENCES

(1) Knoot, C. J.; Ungerer, J.; Wangikar, P. P.; Pakrasi, H. B. Cyanobacteria: Promising biocatalysts for sustainable chemical production. J Biol Chem 2018, 293 (14), 5044–5052. DOI: 10.1074/jbc.R117.815886 From NLM Medline.

(2) Till, P.; Toepel, J.; Buhler, B.; Mach, R. L.; Mach-Aigner, A. R. Regulatory systems for gene expression control in cyanobacteria. Appl Microbiol Biotechnol 2020, 104 (5), 1977–1991. DOI: 10.1007/s00253-019-10344-w From NLM Medline.

(3) Nakahira, Y.; Ogawa, A.; Asano, H.; Oyama, T.; Tozawa, Y. Theophylline-dependent riboswitch as a novel genetic tool for strict regulation of protein expression in Cyanobacterium *Synechococcus elongatus* PCC 7942. Plant Cell Physiol 2013, 54 (10), 1724–1735. DOI: 10.1093/pcp/pct115 From NLM Medline.

(4) Sakamaki, Y.; Maeda, K.; Nimura-Matsune, K.; Chibazakura, T.; Watanabe, S. Characterization of a cyanobacterial rep protein with broad-host range and its utilization for expression vectors. Front Microbiol 2023, 14, 1111979. DOI: 10.3389/fmicb.2023.1111979 From NLM PubMed-not-MEDLINE.

(5) Opel, F.; Siebert, N. A.; Klatt, S.; Tullinghoff, A.; Hantke, J. G.; Toepel, J.; Buhler, B.; Nurnberg, D. J.; Klahn, S. Generation of Synthetic Shuttle Vectors Enabling Modular Genetic Engineering of Cyanobacteria. ACS Synth Biol 2022, 11 (5), 1758–1771. DOI: 10.1021/acssynbio.1c00605 From NLM Medline.

(6) Case, A. E.; Atsumi, S. Cyanobacterial chemical production. J Biotechnol 2016, 231, 106–114. DOI: 10.1016/j.jbiotec.2016.05.023 From NLM Medline.

(7) Li, H.; Shen, C. R.; Huang, C. H.; Sung, L. Y.; Wu, M. Y.; Hu, Y. C. CRISPR-Cas9 for the genome engineering of cyanobacteria and succinate production. Metab Eng 2016, 38, 293–302. DOI: 10.1016/j.ymben.2016.09.006 From NLM Medline.

(8) Li, H.; Pham, N. N.; Shen, C. R.; Chang, C. W.; Tu, Y.; Chang, Y. H.; Tu, J.; Nguyen, M. T. T.; Hu, Y. C. Combinatorial CRISPR Interference Library for Enhancing 2,3-BDO Production and Elucidating Key Genes in Cyanobacteria. Front Bioeng Biotechnol 2022, 10, 913820. DOI: 10.3389/fbioe.2022.913820 From NLM PubMed-not-MEDLINE.

(9) Santos-Merino, M.; Singh, A. K.; Ducat, D. C. New Applications of Synthetic Biology Tools for Cyanobacterial Metabolic Engineering. Front Bioeng Biotechnol 2019, 7, 33. DOI: 10.3389/fbioe.2019.00033 From NLM PubMed-not-MEDLINE.

(10) Jeffrey L. Ried, A. C. An nptI-sacB-sacR cartridge for constructing directed, unmarked mutations in Gram-negative bacteria by marker exchange-eviction mutagenesis. Gene 1987, 57 (2-3), 239–246. DOI: 10.1016/0378-1119(87)90127-2.

(11) Reyrat, J. M., Pelicic, V., Gicquel, B., & Rappuoli, R. Counterselectable markers: untapped tools for bacterial genetics and pathogenesis. Infection and immunity 1998. DOI: 10.1128/IAI.66.9.4011-4017.1998.

(12) Cheah, Y. E.; Albers, S. C.; Peebles, C. A. A novel counter-selection method for markerless genetic modification in *Synechocystis* sp. PCC 6803. Biotechnol Prog 2013, 29 (1), 23–30. DOI: 10.1002/btpr.1661 From NLM Medline.

(13) Nishimoto, K.; Furushima, T.; Kadowaki, T.; Jimbo, H.; Yamamoto, N.; Ikeda, K.; Takatani, N.; Aichi, M.; Omata, T.; Nishiyama, Y. Overexpression of endogenous galactolipases and an efflux transporter enhances the secretion of extracellular free fatty acids by *Synechococcus elongatus* PCC 7942. Biotechnol Biofuels Bioprod 2026, 19 (1). DOI: 10.1186/s13068-026-02768-0 From NLM PubMed-not-MEDLINE.

(14) Niu, T. C.; Lin, G. M.; Xie, L. R.; Wang, Z. Q.; Xing, W. Y.; Zhang, J. Y.; Zhang, C. C. Expanding the Potential of CRISPR-Cpf1-Based Genome Editing Technology in the Cyanobacterium *Anabaena* PCC 7120. ACS Synth Biol 2019, 8 (1), 170–180. DOI: 10.1021/acssynbio.8b00437 From NLM Medline.

(15) Hasegawa, H.; Kanesaki, Y.; Watanabe, S.; Tanaka, K. A high-temperature sensitivity of *Synechococcus elongatus* PCC 7942 due to a tRNA-Leu mutation. J Gen Appl Microbiol 2023, 69 (3), 167–174. DOI: 10.2323/jgam.2023.02.001 From NLM Medline.

(16) Nakano, C.; Kim, H. K.; Ohnishi, Y. Identification of the first bacterial monoterpene cyclase, a 1,8-cineole synthase, that catalyzes the direct conversion of geranyl diphosphate. Chembiochem 2011, 12 (13), 1988–1991. DOI: 10.1002/cbic.201100330 From NLM Medline.

(17) Sakamaki, Y.; Ono, M.; Shigenari, N.; Chibazakura, T.; Shimomura, K.; Watanabe, S. Photosynthetic 1,8-cineole production using cyanobacteria. Biosci Biotechnol Biochem 2023, 87 (5), 563–568. DOI: 10.1093/bbb/zbad012 From NLM Medline.

(18) Juergens, U. R.; Dethlefsen, U.; Steinkamp, G.; Gillissen, A.; Repges, R.; Vetter, H. Anti-inflammatory activity of 1.8-cineol (eucalyptol) in bronchial asthma: a double-blind placebo-controlled trial. Respir Med 2003, 97 (3), 250–256. DOI: 10.1053/rmed.2003.1432 From NLM Medline.

(19) Lejars, M.; Hajnsdorf, E. RNase III Participates in the Adaptation to Temperature Shock and Oxidative Stress in *Escherichia coli*. Microorganisms 2022, 10 (4). DOI: 10.3390/microorganisms10040699 From NLM PubMed-not-MEDLINE.

(20) Ohdate, K.; Sakata, M.; Maeda, K.; Sakamaki, Y.; Nimura-Matsune, K.; Ohbayashi, R.; Hess, W. R.; Watanabe, S. Discovery of novel replication proteins for large plasmids in cyanobacteria and their potential applications in genetic engineering. Front Microbiol 2024, 15, 1311290. DOI: 10.3389/fmicb.2024.1311290 From NLM PubMed-not-MEDLINE.

(21) Watanabe, S.; Ohbayashi, R.; Shiwa, Y.; Noda, A.; Kanesaki, Y.; Chibazakura, T.; Yoshikawa, H. Light-dependent and asynchronous replication of cyanobacterial multi-copy chromosomes. Mol Microbiol 2012, 83 (4), 856–865. DOI: 10.1111/j.1365-2958.2012.07971.x From NLM Medline.

