## Supplemental data for "Development of a Temperature-Based Method for Repeated Markerless Gene Integration in cyanobacterium *Synechococcus elongatus* PCC 7942"

### 1. Table

**Table S1. Strain**

| Name | Genotype | references |
|---|---|---|
| WT | Wild type strain | (21) |
| H1 | Temperature sensitive (TS) phenotype, tRNA-Leu G60T | (15) |
| A | tRNA-Leu G60T, Δ*nynA*:: *aad* | this study |
| G | tRNA-Leu G60T, Δ*nynA*:: *gfp* | this study |
| GN | tRNA-Leu G60T, Δ*nynA*:: *gfp*, NS85:: *nynA-cat* | this study |
| GS | tRNA-Leu G60T, Δ*nynA*:: *gfp*, NS85:: $P_{cpcB560}$-*mScarlet* | this study |
| GSN | tRNA-Leu G60T, Δ*nynA*:: *gfp*, NS85:: $P_{cpcB560}$–*mScarlet,* NS98:: *aad-nynA* | this study |
| GSL | tRNA-Leu G60T, Δ*nynA*:: *gfp*, NS85:: $P_{cpcB560}$–*mScarlet,* NS98:: $P_{trc}$-riboswitch-*luciferase* | this study |
| C | tRNA-Leu G60T, Δ*nynA*:: *lacI*–$P_{trc}$–*cnsA*$^{opt}$ | this study |

**Table S2. Plasmid**

| Name | Features | References |
|---|---|---|
| p1640_aad | p1640, *nynA* upstream region (1707135-1707880), *aad*, *nynA* downstream region (1708415-1709165) | this study |
| p1640_gfp | p1640, *gfp* gene | this study |
| pNS85_nynA | pUC19 (*bla*, *colE1*), neutral site 85 upstream region (83869-82712), *nynA* gene (1707682-1708414), *cat*, neutral site II downstream region (82711-80756) | this study |
| pNS85_mScarlet | pUC19 (*bla*, *colE1*), neutral site 85 upstream region (83869-82712), $P_{cpcB560}$-*mScarlet*, neutral site 85 downstream region (82711-80756) | this study |
| pNS98_nynA | pBR322 (*bla*, *colE1*), neutral site 98 upstream region (2578672-2579525), *aad*, *nynA* gene (1707682-1708414), neutral site 98 downstream region (2579566-2580405) | this study |
| pNS98_luciferase | pBR322 (*bla*, c*olE1*), neutral site 98 upstream region (2578672-2579525), $P_{trc}$-riboswitch-*luciferase*, neutral site 98 downstream region (2579566-2580405) | this study |
| p1640_cnsA | p1640, *lacI*, $P_{trc}$-$cnsA^{opt}$ geme | this study |

**Table S3. Oligonucleotide primers used in this study.**

| Primer | Sequence (5′ to 3′) [a] |
|---|---|
| *Primers for plasmid construction* | |
| F1 | AACGACGGCCAGTGAGTGCGATCGCTCAGTTTTGGCT |
| R2 | AAACAGCTATGACCATTAGGGAAGAACCTCAAGGCTGGCCGCT |
| R3 | TCACTGGCCGTCGTTTTACA |
| F4 | TGGTCATAGCTGTTTCCTGT |
| F5 | CCACTCCTCCAGATCCGGGATCTCGACGCTCTCCCTTATG |
| R6 | TTCGTTTCTCAGGGATTATTTGTATAGTTCATCCATGCCATGT |
| F7 | CCACTCCTCCAGATCCCAGGCATCAAATAAAACGAAAGGCTCAGTCG |
| R8 | TTCGTTTCTCAGGGACACATTTCCCCGAAAAGTGCCACCTGACGTC |
| F9 | TCCCTGAGAAACGAATAATTTTCAGC |
| R10 | GATCTGGAGGAGTGGCACTTACTTCGT |
| F11 | AGTTTGTGGGTCATTTCGAAGCCGCCGATGGGGCTT |
| R12 | GGATTTATTTATTCTTTAAGACTGGCGCCGCCAATGTTCCAG |
| F13 | AGTTTGTGGGTCATTACCTGTAGAGAAGAGTCCCTGAA |
| R14 | GACAGTCAGCCATGGAAGAGTTTGTAGAAACGCAAAAAGGC |
| F15 | AGAATAAATAAATCCTGGTGTCCCTGTTG |
| R16 | AATGACCCACAAACTGCTCAAACAT |
| F17 | CCATGGCTGACTGTCTCAAACTAA |
| F18 | GTTACCACCGCTGCGTCGAAGCCGCCGATGGGGCT |
| R19 | GACTCCACGTCGATATTAAGACTGGCGCCGCCAATGTTCCAG |
| F20 | CTTTCTGCGTTTATA CAAATATTCTGAAATGAGCTGTTGAC |
| R21 | GACCGCTTCTGCGTTCTG |
| F22 | TATCGACGTGGAGTCGATCACTGTGA |
| R23 | CGCAGCGGTGGTAACGGCGC |
| F24 | AACGCAGAAGCGGTCTATCGACGTGGAGTCGATCA |
| R25 | TATAAACGCAGAAAGGCTTCTGGAGCAGGAAGATGT |
| *Primers for segregation analysis* | |
| F26 | CCGGCAGGATCAACCGCAGAGGATTGTGAC |
| R27 | GCACTACGGCCAGCGGTGGGCG |
| F28 | ATAGCGGAACGCATCACTAATCGCG |
| R29 | GCGATACGGGTACGGCGTTGATAGT |
| F30 | CAAACAGGTGCAGCAGCAACT |
| R31 | CATCGCTATCTCTTAGGACTTCGCAG |

### 2. Figure

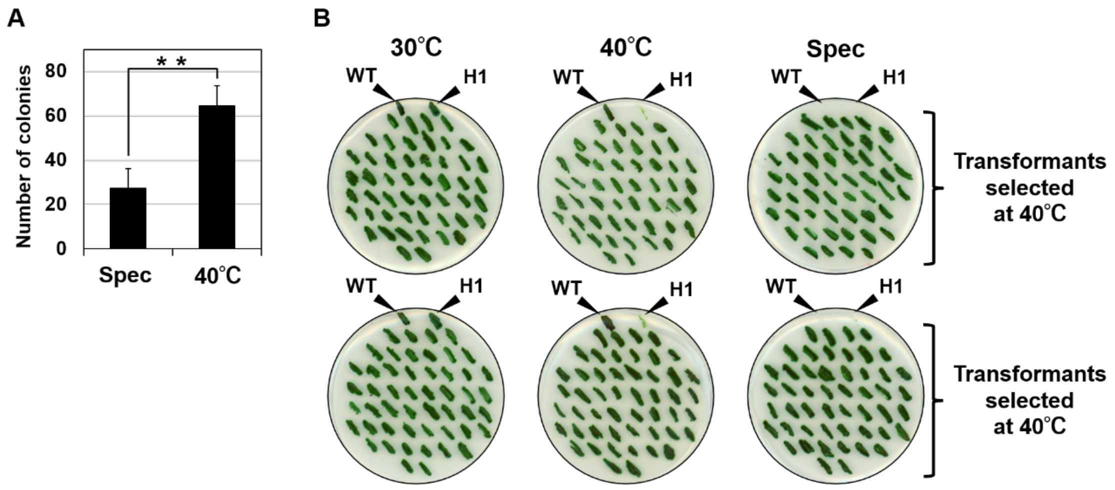


**Figure S1. Evaluation of high-temperature selection in the H1 strain.**

(A) Comparison of the numbers of colonies obtained by selection with spectinomycin (Spec) and by incubation at 40 °C. The number of colonies obtained under each selection condition was determined. Data is presented as the mean ± SD of five independent experiments (n = 5), ** $P < 0.01$ by paired Student's *t*-test. (B) Complete plate images showing all 100 transformants tested for temperature and spectinomycin resistance. A close-up representative section of these images is shown in Fig. 2C.

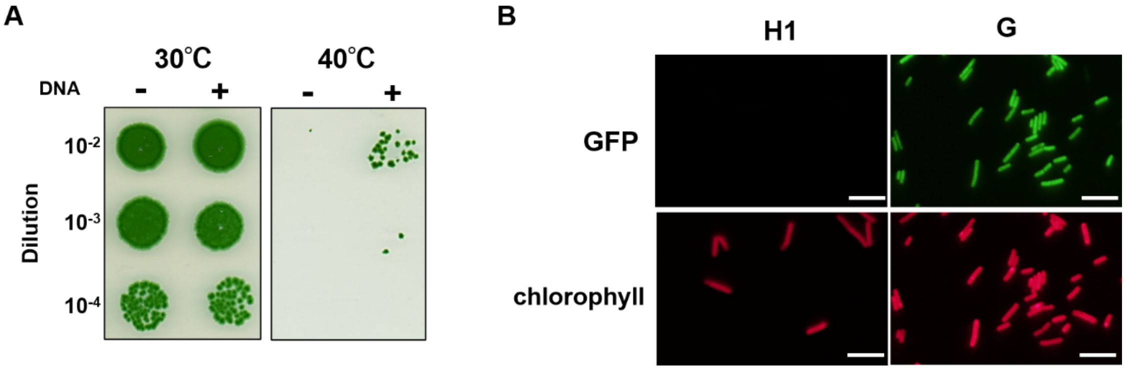


**Figure S2. Selection and fluorescence characterization of the *gfp*-expressing strain.**

(A) Selection of transformants by incubation at 40 °C. Transformants were diluted and spotted onto BG-11 agar plate and incubated on antibiotic-free BG-11 agar plates at 40 °C for 8 days. (B) Fluorescence microscopy images of the H1 and G strains showing GFP fluorescence and chlorophyll autofluorescence. Scale bar, 10 µm.

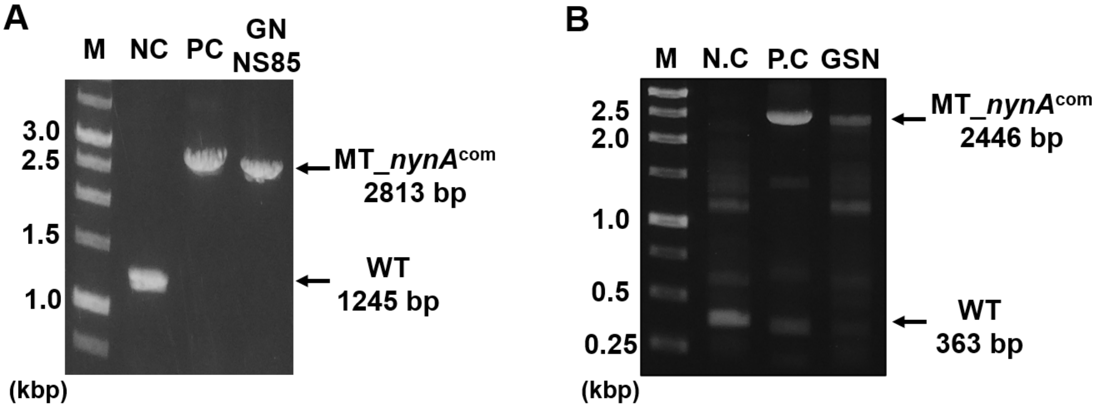


**Figure S3. PCR verification of nynA complementation at NS85 and NS98.**

(A) PCR analysis of the G and GN strains. Primers shown as green arrows in Fig. 4A were used to confirm integration of *nynA* at the NS85 locus. M, DNA marker; NC, G strain; PC, pNS85_nynA plasmid (Table S1). (B) PCR analysis of the GS and GSN strains. Primers shown as orange arrows in Fig. 6A were used to confirm integration of *nynA* at the NS98 locus. M, DNA marker; NC, GS strain; PC, pNS98_nynA plasmid (Table S1).

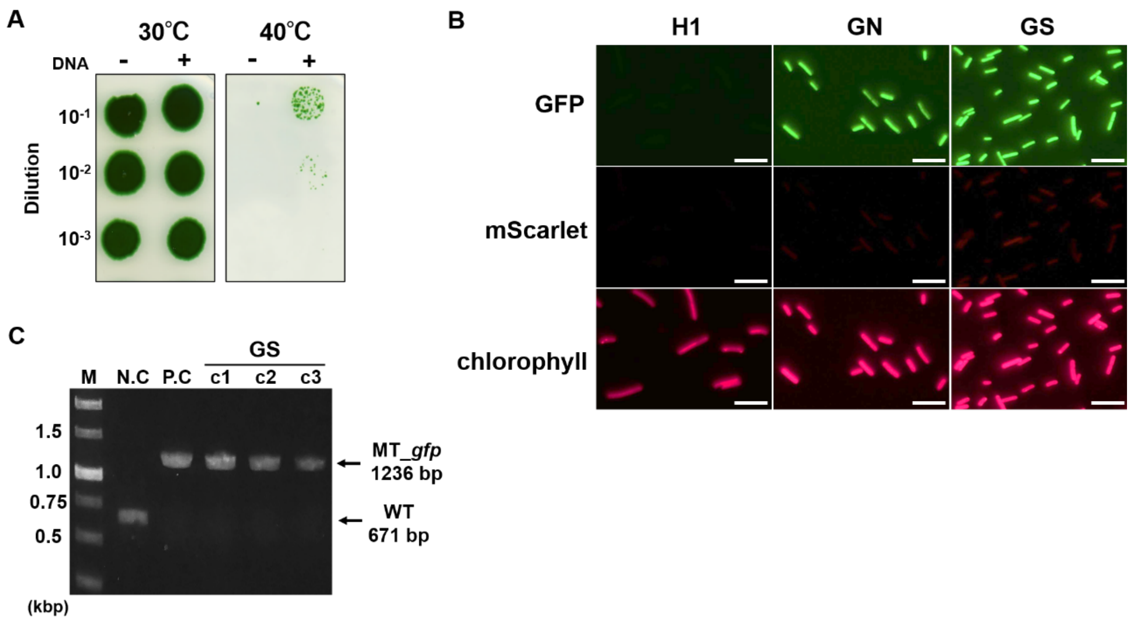


**Figure S4. Selection and fluorescence characterization of the *mScarlet*-expressing strain.**

(A) Selection of transformants by incubation at 40 °C. Transformants were selected on antibiotic-free BG-11 agar plates at 40 °C for 8 days. (B) Fluorescence microscopy images of the H1, GN and GS strains showing GFP fluorescence, mScarlet fluorescence and chlorophyll autofluorescence. Scale bar, 10 µm. (C) PCR analysis of the GN strain and three independent transformants (c1–c3). Primers shown as green arrows in Fig. 3A were used to confirm the presence of the *gfp* gene. M, DNA marker; NC, GS strain; PC, p1640_gfp plasmid (Table S1).

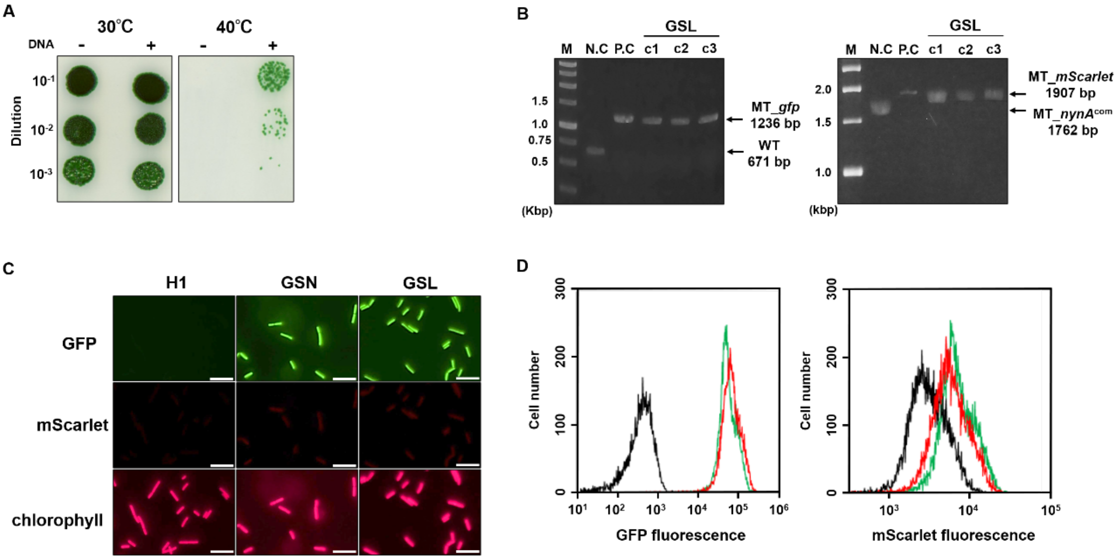


**Figure S5. Selection and fluorescence characterization of the *luciferase*-expressing strain.**

(A) Selection of transformants by incubation at 40 °C. Transformants were selected on antibiotic-free BG-11 agar plates at 40 °C for 9 days. (B) PCR analysis of the GSN strain and three independent transformants (c1–c3). Primers shown as green arrows in Fig. 3A were used to confirm the presence of the *gfp* gene, and primers shown as orange arrows in Fig. 5A were used to confirm the presence of the *mScarlet* gene. M, DNA marker; NC, GSN strain; PC, p1640_gfp plasmid for *gfp* detection and pNS85_mScarlet plasmid for *mScarlet* detection (Table S1). PCR products for *mScarlet* detection were separated on a 1.8% agarose gel in TBE buffer at 50 V. (C) Fluorescence microscopy images of the H1, GSN, and GSL strains showing GFP fluorescence, mScarlet fluorescence, and chlorophyll autofluorescence. Scale bar, 10 µm. (D) Flow cytometric analysis of GFP and mScarlet fluorescence. Histograms of GFP and mScarlet fluorescence are shown for the H1 strain (black), GSN strain (red), and GSL strain (green).
